# Gaussian Network Model Revisited: Effects of Mutation and Ligand Binding on Protein Behavior

**DOI:** 10.1101/2021.09.29.462446

**Authors:** Burak Erman

## Abstract

The coarse-grained Gaussian Network model, GNM, considers only the alpha carbons of the folded protein. Therefore it is not directly applicable to the study of mutation or ligand binding problems where atomic detail is required. This shortcoming is improved by including the local effect of heavy atoms in the neighborhood of each alpha carbon into the Kirchoff Adjacency Matrix. The presence of other atoms in the coordination shell of each alpha carbon diminishes the magnitude of fluctuations of that alpha carbon. But more importantly, it changes the graph-like connectivity structure, i.e., the Kirchoff Adjacency Matrix of the whole system which introduces amino acid specific and position specific information into the classical coarse-grained GNM which was originally modelled in analogy with phantom network theory of rubber elasticity. With this modification, it is now possible to make predictions on the effects of mutation and ligand binding on residue fluctuations and their pair-correlations. We refer to the new model as ‘all-atom GNM’. Using examples from published data we show that the all-atom GNM applied to in silico mutated proteins and to their laboratory mutated structures gives similar results. Thus, loss and gain of correlations, which may be related to loss and gain of function, may be studied by using simple in silico mutations only.

## 1. Introduction

Thermal fluctuations of atoms of a folded protein constitute the signature of that protein. They are unique in the unperturbed state but tend to change upon perturbation such as mutation of a residue or binding of a ligand, both of which change the degree of local packing in the system. Fluctuations of an atom in condensed matter are directly affected by the presence of its spatial neighbors. A large number of neighbors results in smaller fluctuations as may be seen from numerous crystal structures in the Protein Data Bank^1^. This inverse relationship between the number of neighbors and fluctuations forms the basis of the classical Gaussian Network Model (GNM) of proteins^2^. The GNM is a coarse-grained model where only the alpha carbons are considered. The number of alpha carbons within a sphere of a given cutoff distance from a residue is designated as the number of neighbors of that residue. The coarse-grained GNM is constructed^2^ in complete analogy with the theory of elastomeric phantom networks^3^ where an alpha carbon corresponds to a network junction. A junction in a phantom network is connected to its neighbors with linear springs in the same way as an alpha carbon is connected to its spatial neighbors with fictitious springs. The space surrounding a given junction in a phantom network is filled with atoms^4^ which do not contribute to the elasticity of the network. Only those junctions covalently connected to the network contribute. Likewise, in the GNM, only spatially neighboring alpha carbons contribute to the elasticity of the protein^2^. In more detail, an f-functional junction in an elastic phantom network is equivalent to an alpha carbon with f other alpha carbons within its first coordination shell. The first coordination shell is defined as a sphere centered around the given alpha carbon and with a radius of a given cutoff value. Within the GNM analogy, the mean-square fluctuations, ⟨(Δ*R*_*i*_)^2^⟩, of a residue i, assumed to be collapsed on its alpha carbon may be written as^3, 5^

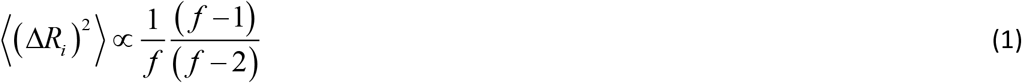

where, *f* is the number of neighboring alpha carbons within a cutoff distance of residue i. A phantom network and the protein are both represented by a graph connectivity matrix, also referred to as the Kirchoff adjacency matrix^6^, Γ (See the Models and Methods section for detailed definition), whose inverse is proportional to the correlation of fluctuations of residue pairs. We present the equations of the GNM in a historical perspective in the theory section. Although the elements of Γ are obtained by considering the spatial neighbors of residues only, the inverse operation on Γ introduces the effects of interactions between spatially distant residue pairs into the formulation. In this sense, the model is capable of handling allosteric interactions. The GNM rests on a simple basic feature of proteins: The number of neighbors of each residue and the relationship of this number to the mean-square fluctuations of a residue which is approximated by Eq. 1. Interestingly, since the Γ matrix is obtained from the contact map of the protein, and since it is possible to approximate the three dimensional structure of a protein from its contact map^7^, the GNM is essentially a model that implicitly relates shape and structure to function of proteins, where function is visualized in terms of correlation of fluctuations. If two alpha carbons fluctuate in full harmony, always in the same direction, then the two are said to be positively correlated. If they always move in opposite directions, then they are anti-correlated. If their fluctuations are random relative to each other, then they are uncorrelated. A protein performs its function as a result of information transmitted from one point to the other in the protein. Information can be transmitted from one residue to another only if the fluctuations of the two points are correlated. Thus, correlations play a significant role in the function of the protein. Understanding the changes in correlations upon mutation or ligand binding is even more important because they reflect the changes in the function of the protein: loss of correlations may be associated with loss of function. Likewise, production of new correlations may be associated with gain of function.

Although the GNM is a coarse-grained model, it led to results that could explain and predict several aspects of proteins. However, the phantom like nature of the GNM implies that the effects of local atomic packing, a strong function of (i) residue type, and (ii) position in the protein^8^ are missing. The local environment of a given residue within a cutoff distance of 4.0 Å, for example, contains between 20 and 100 atoms, including the hydrogen atoms. This variance in local packing density, whose effect is not built into the GNM, is too large to be ignored in a realistic protein model. The fluctuations of atoms in a protein are large, in the order of several Ångstroms. Obviously, the contacting atoms from the neighboring amino acids greatly restrict the fluctuations of a given atom. The important factor is the dependence of these restrictions on residue type and position in the protein. Atom-atom contacts obtain values that minimize the contact energy of a given residue and its neighbors by minimizing the free space around the given residue, as in a jigsaw puzzle, subject to the constraints imposed by the tertiary structure of the protein. The geometry of the specific residue plays significant role in this minimization and jigsaw puzzle-like rearrangement. Trying to fit a large residue into a small cavity in the contacting region results in large repulsive forces which distort the chain, which in itself has a large energy cost. Furthermore, at the atomic level, different residues may form contacts to atoms that lead to different paths in the contact matrix. This is the topological dispersion where the contacts of one residue type may lead to interactions in a certain region of the protein (or the graph) while another residue type may lead to interactions at a totally different region. Therefore, replacement of an amino acid with another may have significant nonlocal effects in the protein. In this respect, local packing density has far-reaching consequences in protein behavior. It has been shown that local packing density is the main determinant of the rate of protein sequence evolution^9^ which affects the stability and dynamics,^10^ which relate to fluctuations, and hence may lead to different long-range excitations. Inasmuch as effects carry to large distances away from the residue considered, their effects will be seen most strongly in the slow modes of the protein. There has been an earlier attempt to improve the coarse-grained GNM by considering four heavy atoms *C*^*α*^, *C, N, C*^*β*^ and their combinations, extracting energy functions from known crystal structures^11^, modifying the coarse graining accordingly and comparing the resulting B-factors with those of the coarse-grained GNM. The present model focusses on the packing density around each atom and its dependence on mutation and ligand binding, counts every atom, and emphasizes changes in residue pair-correlations, hence differs from the previous study.

The constraining effect of the atoms around a given junction other than its covalently bonded neighbors in a rubber or elastomer was evident in stretching or swelling it. Stretching or swelling decreased the local packing density of atoms around a given junction. The effects of the decrease of packing density turned out to be of utmost importance in the elasticity of rubber. When the effects of constraints were taken into account in an improved model known as the Flory-Erman theory, almost all deviations of the simple phantom network model from experiment were accounted for^4, 5, 12-16^. In the protein, the counterparts of strain and swelling of rubber elasticity are the mutation of a residue, or binding of a ligand, both of which affect the local packing density in the protein. Changing the local packing around a given residue may change (i) the mean positions of atoms, or (ii) the fluctuations of atoms in the protein without change in mean positons. The first one is an energetically expensive operation that will lead to correlated spatial rearrangement of mean positions, i.e., the shape of the protein. The model that we propose can treat case (ii). Finding the new mean positions of the perturbed protein requires extensive molecular dynamics simulations or experimental structure determination of the perturbed system. The experimental examples we present in this paper show that conformational change upon mutation or ligand binding is not significant and case (ii) may be a good approximation. More specifically, the model will be sufficiently rigorous for proteins that exhibit dynamic allostery, i.e., allostery without conformational change, where a perturbation, mutation or ligand binding results in correlations between distant residues^17^.

## 2. Model and methods

### i. Difference between coarse-grained GNM and all-atom GNM

The packing of atoms in a typical protein is shown in Figure 1. The alpha carbon of ILE359 of the PDZ domain, 1BE9.pdb is shown as the central gray sphere. The alpha carbons of the neighboring residues within a given cutoff distance, of ILE359, for example, are shown as black spheres. The white spheres are the heavy atoms affixed to the amino acids whose alpha carbons are shown. Hydrogen atoms are not shown. In coarse-grained GNM, the presence of the white spheres are ignored and the black spheres are attached to the central gray sphere with linear springs. For the whole protein, every pair that are within the coordination shell of each other is connected by such springs. In this respect the model is identical to the phantom network model of rubber elasticity where alpha carbons are replaced by junctions or cross-links, and springs are between covalently bonded junctions. Each alpha carbon fluctuates about its mean position. Larger number of alpha carbons in the neighborhood of a given alpha carbon results in a smaller amplitude for its fluctuations. In the real picture of the protein, as in the real elastomer, several other atoms fill the space in the coordination shell of the chosen reference atom. The presence of each atom decreases the fluctuations of the alpha carbons. In rubber elasticity jargon^4, 9, 11-1^ the action of these additional atoms are ‘the constraints acting on the fluctuations of the given junction’. Thus, as in the coarse-grained GNM, it is the number of neighboring atoms that matters. The contribution of the all-atom GNM lies in the fact that the identity of each amino acid is introduced into the Γ matrix which did not exist in previous theories.

**Figure 1.**
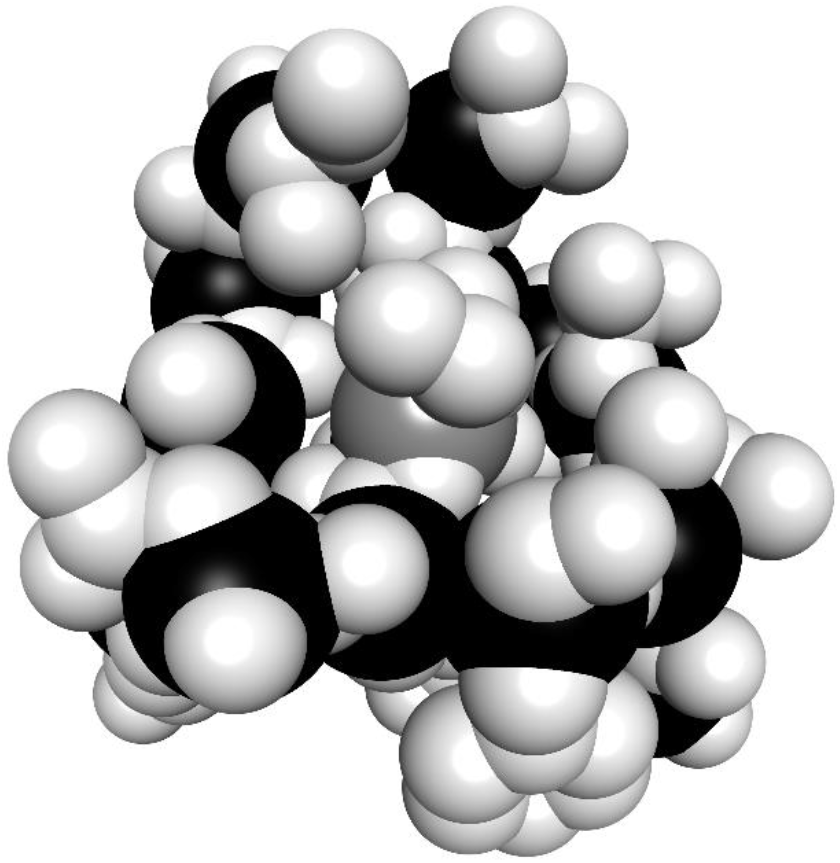
ILE359 (gray sphere) and its first neighbors. Black spheres are the neighboring alpha carbons and the white spheres are the heavy atoms other than alpha carbons.

The statistical mechanics of phantom networks was developed by James and Guth in 1943^18^. The constraining effects of the atoms on the fluctuations of a junction was first realized 32 years later by Ronca and Allegra in 1975^19^, and by P. J. Flory in 1976^3^ which eventually led to the constraint theories of rubber elasticity. Likewise, the coarse-grained GNM was modelled after the elastic phantom network model by the present author and his collaborators in 1997, and the improvement by including the effects of constraints, like that of constraint theories of rubber elasticity, is the next step which is introduced in this paper.

In Figure 1, the reference alpha carbon ILE359 has 14 other alpha carbons in its first coordination shell whose radius we take as 7.2 Å ^20^. In addition to the 14 alpha carbons, there are 96 other heavy atoms. Their presence decreases the fluctuations of ILE359. If ILE359 is mutated into another amino acid the number of atoms and therefore the packing density within the coordination shell will change. The all-atom GNM presented here accounts for the effects of these changes on fluctuations and correlations in the system.

### ii. The Γ matrix and correlations of fluctuations

The Kirchoff adjacency or simply the connectivity matrix is now defined as:

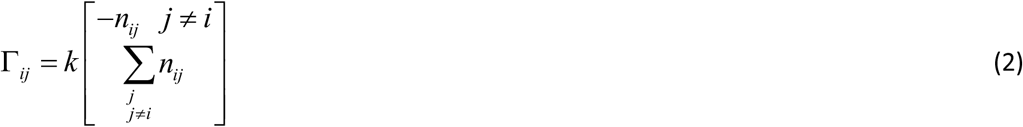

Where i and j represent the residue indices, *n*_*ij*_ is the number of atom pairs closer than a given cutoff distance, one belonging to residue i the other to residue j. k is a proportionality factor. There are two differences between the Γ matrices of the coarse-grained and all-atom GNM: (i) The magnitudes of the entries change, but more importantly (ii) new terms in the all-atom Γ will appear because of atom-atom contacts in addition to the alpha carbon contacts in the coarse-grained Γ. This change in the Γ matrix will reflect the change in the graph structure of the system and may lead to significant changes in protein behavior. A point of importance which we do not elaborate in the present paper is the change in the dynamics of the system by mutation or ligand binding. The use of the Γ matrix in the equation of motion, i.e.,the Langevin equation and its solution are given in detail in a recent paper^21^. The correlation of the fluctuation of two residues is expressed as

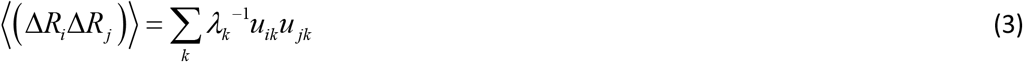

where, the angular brackets denote time or ensemble average, Δ*R*_*i*_ is the fluctuation of residue i from its mean position in the protein, λ_*k*_ is the kth eigenvalue of the Γ matrix, and *u*_*ik*_ is the ith element of the eigenvector corresponding to the kth eigenvalue. In this sense the only difference of the all-atom GNM from the coarse-grained GNM is in the derivation of the Γ matrix. As in the coarse-grained GNM, the problem is isotropic in which different coordinate directions are ignored which amounts to considering all coordination shells, i.e., the one shown in Figure 1 as isotropic and spherically symmetrical.

Rendering the fluctuation correlation expression, Eq. 3, in terms of eigenvalues and eigenvectors gives the possibility of studying mutation effects in different length scales. All results on correlations reported in this paper are obtained using all modes of the protein.

### iii. Scaling the results and the correlation matrix

The all-atom GNM, like the coarse-grained GNM contains a single constant that has to be adjusted to match known results. Experimental B-factors have been used as reference in previous work^2^. The interest in this paper is on the changes in pair correlations. For this purpose, we define the normalized correlation, *C*_*i, j*_ as

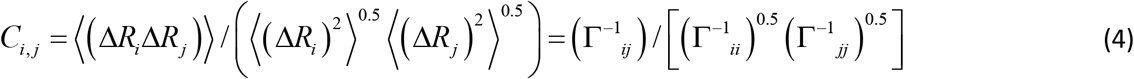

In this form, the constant needed to adjust the calculation results to experimental data cancels out. Elimination of the adjustable parameter of the model by using normalized correlations is an important advantage.

### iv. In silico mutation of the proteins

Mutations of amino acids are performed using Biovia Discovery Studio Visualizer^22^. After each mutation, the amino acid is selected and its energy is minimized, keeping the rest of the protein fixed, using the Dreiding force-field. This minimization modifies the bond lengths and bond angles of the mutated amino acid such that its energy plus the energy of interaction of its atoms with their neighbors is minimized.

This operation changes the atomic packing around the mutated amino acid, which is the only source of new information in the Γ matrix. Most of the atom-atom interactions around the mutated amino acid are within 4 Ångstroms, therefore the cutoff distance for the Γ matrix is chosen as 4 Å in contrast to 7.2 Å for the coarse-grained GNM.

In reality, mutating one amino acid into another may affect the atom positions of neighboring amino acids, which may further result in the change of conformation of the full protein. The performance of the all-atom GNM needs be checked on actual laboratory mutated samples too. For this purpose, we apply the all-atom GNM also to structures whose crystal structures are determined after mutation. As will be discussed below, the results for in silico mutated and laboratory mutated samples do not show significant difference.

## 3. Results and Discussion

Here, we test the predictions of the all-atom GNM by comparing with experimental and computational data from the literature. The proof-of-concept case that we report here, allostery in a PDZ domain, is used mainly for two purposes: (i) to show what all-atom GNM is able to do but coarse-grained GNM is not, and (ii) to show that all-atom GNM applied to in silico mutated structures and laboratory mutated crystal structures gives similar results.

In a recent study, ligand binding and mutation effects in the PDZ domain were studied experimentally^23^ in which crystal structures of wild type and mutated proteins were given.

PDZ domains are 100-amino acid mixed a/b folds that recognize the C-terminal region of target proteins at the active site groove between the b2 strand and a2 helix. See Figure 1A in Reference ^23^. The binding capability of the ligand CRIPT depends on the mutation G330T. Here, we use all-atom GNM to study the effects resulting from this mutation. In particular, we compare the effects of mutation on ligand binding with the effects of ligand binding on the mutated protein.

### i. Effect of mutation G330T on ligand bound protein

The wild type structure that contains the ligand CRIPT is 1BE9.pdb. The G330T mutation is applied to 1BE9 using Biovia Discovery Studio Visualizer as described above and the correlation differences of residue 330 with all other residues of the protein are calculated with all-atom GNM. These differences are shown by the thick black curve in Figure 2.

**Figure 2.**
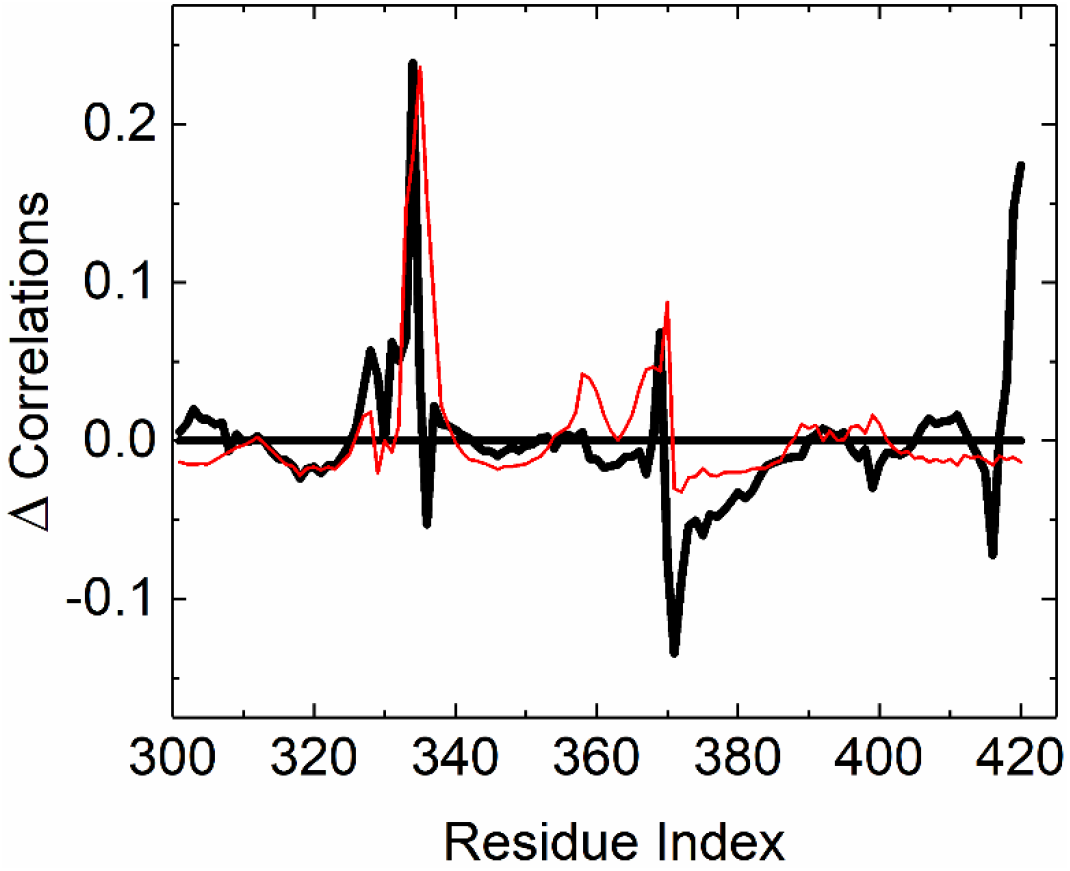
Effect of mutation G330T in the ligand bound protein. Black solid line from in silico mutation of the ligand bound protein, red thin line from laboratory mutated protein. Ordinate values show the differences of correlation of residue 330 with other residues.

Experimentally^23^, the wild type with CRIPT and the G330T structures are available as 5HEB.pdb and 5HEY.pdb, respectively. All-atom GNM is applied to these two structures also, and correlation differences of residue 330 with all other residues of the protein are calculated. Mutation of 330 results in a gain of correlation with 328 and 331, but most importantly a significant gain of correlation with 334 and 370 and a significant loss of correlation with 371. GLU334, ALA370 and SER371 all flank residue 330. There is also a strong correlation gain of 330 and the c-terminal residues.

### ii. Effect of mutation G330T on protein without ligand

The ligand CRIPT is removed from the crystal structure and G330T mutation is applied to 1BE9 using Biovia Discovery Studio Visualizer. The correlation differences of residue 330 with all other residues of the protein are calculated with all-atom GNM. These differences are shown by the thick black curve in Figure 3. Experimentally^23^, the G330T crystal structure without CRIPT is available as 5HET.pdb. All-atom GNM is applied to 5HET also as explained in the previous example. The crystal structure result is shown by the thin red line. Differences in Figure 3 from Figure 2 show the correlation differences due to the absence CRIPT. One important observation is that 330 has now lost correlation with the C-terminal residues.

**Figure 3.**
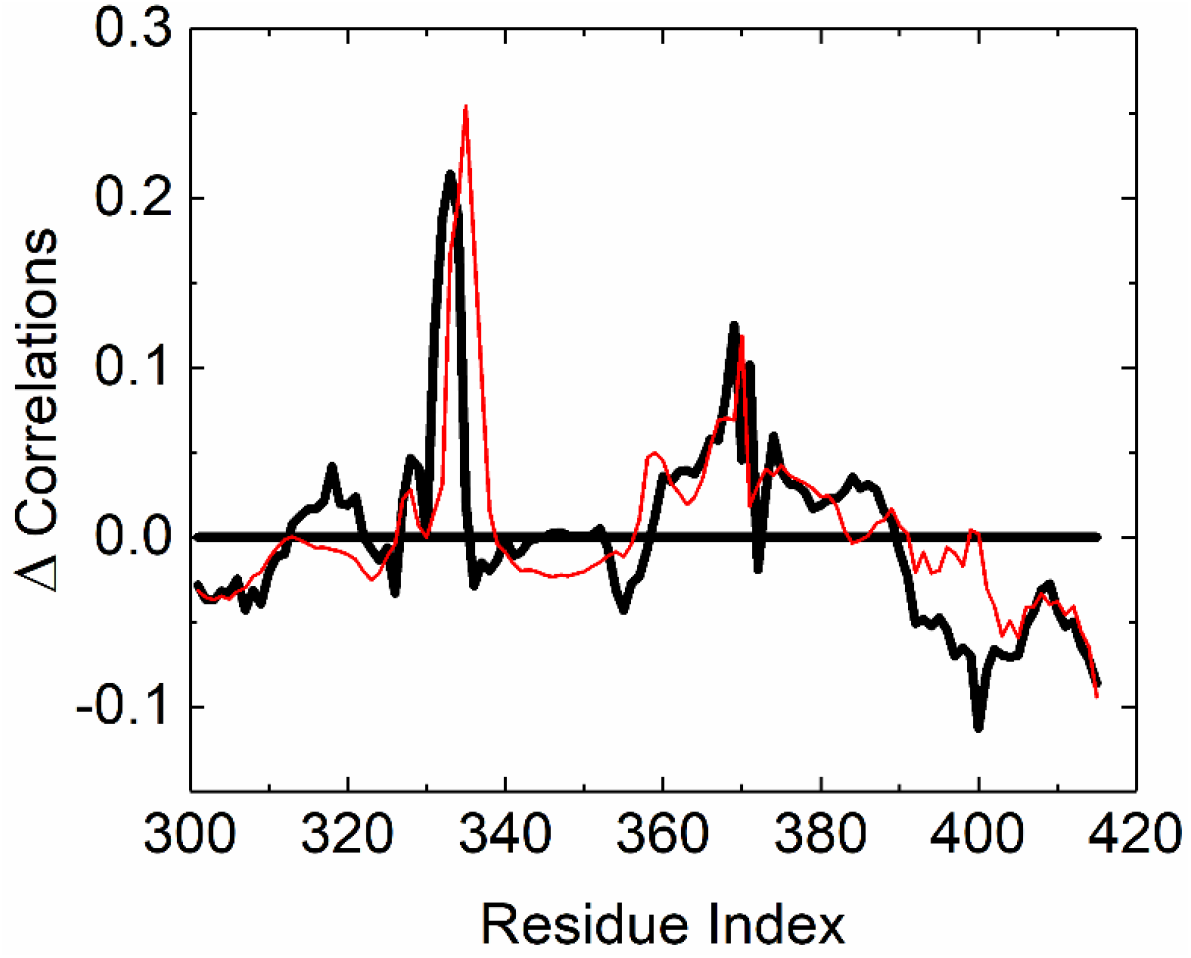
Effect of mutation G330T in the absence of ligand. Black solid line from in silico mutation, red thin line from laboratory mutated proteins. Ordinate values show the differences of correlation of residue 330 with other residues of the protein

### iii. Effect of H372A mutation in ligand bound protein

The *H372A* mutation is applied to 1BE9 using Biovia Discovery Studio Visualizer. Experimentally^23^, the wild type protein with CRIPT and the H372A structures are available as 5HEB.pdb and 5HFB.pdb. All-atom GNM is applied to these structures, and correlation differences of residue 372 with all other residues of the protein are calculated and presented in Figure 4 as the thick black line for the in silico mutation and laboratory mutation as thin red line. Strong loss of correlation of 372 with 330 is observed. Also, 372 exhibits a strong loss of correlation with the C-terminal residues.

**Figure 4.**
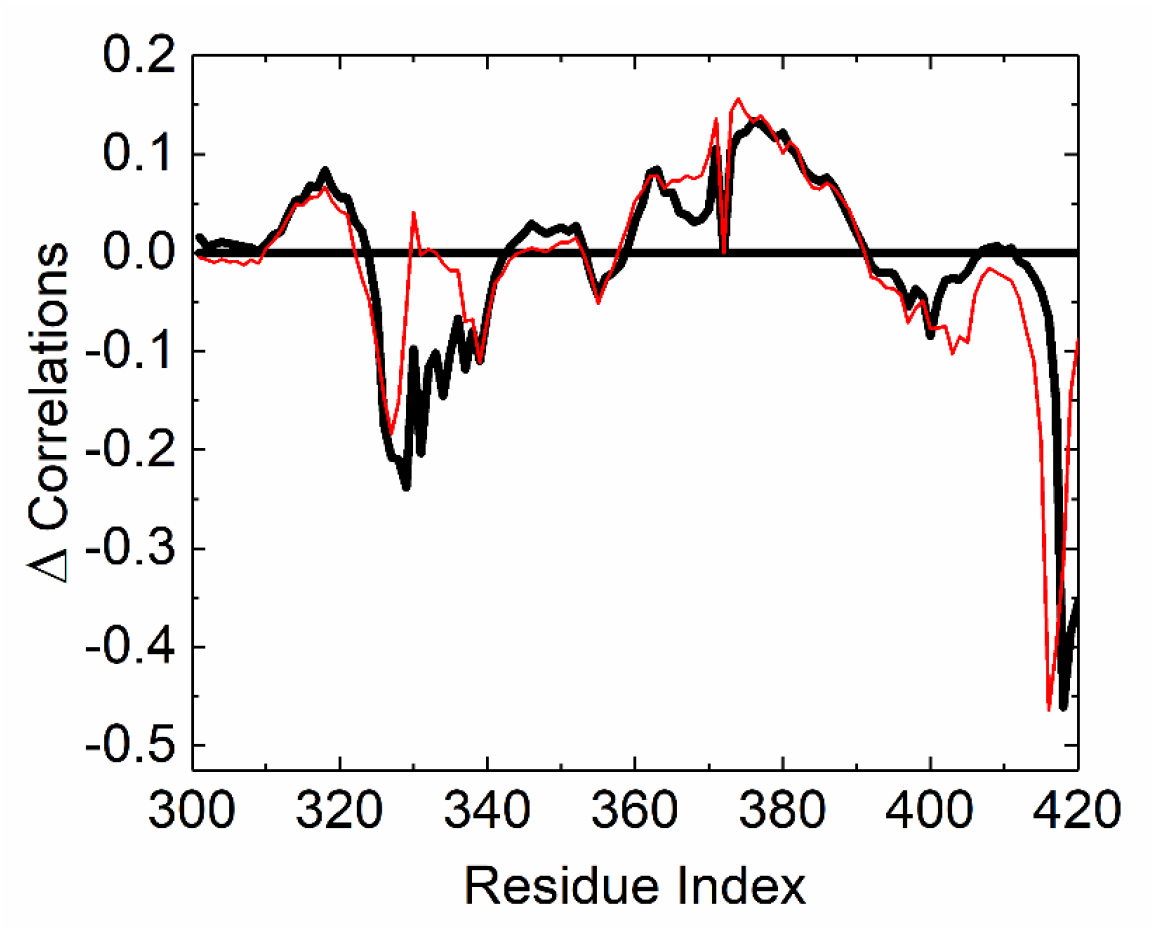
Effect of mutation H372A on the ligand bound protein. Black solid line from in silico mutation, red thin line from laboratory mutation. Ordinate values show the differences of correlation of residue 330 with other residues of the protein.

### iv. Differences between ligand binding on wild type and G330T mutated proteins

There is no experimental structure to study the effect of ligand binding on the apo wild type structure. However, we can now use 1BE9 and an apo structure (where CRIPT is removed from 1BE9) and apply all-atom GNM to determine correlation differences. Similarly we use G330T mutated 1BE9 and an apo structure (where CRIPT is removed from 1BE9 and G330T mutation applied). In Figure 5 we present the correlation differences of residue 330 for ligand binding to mutated and ligand binding to wild type structures. Correlation loss for residue 372 and gain for the c-terminal when ligand binds to mutated structure is worth noting.

**Figure 5.**
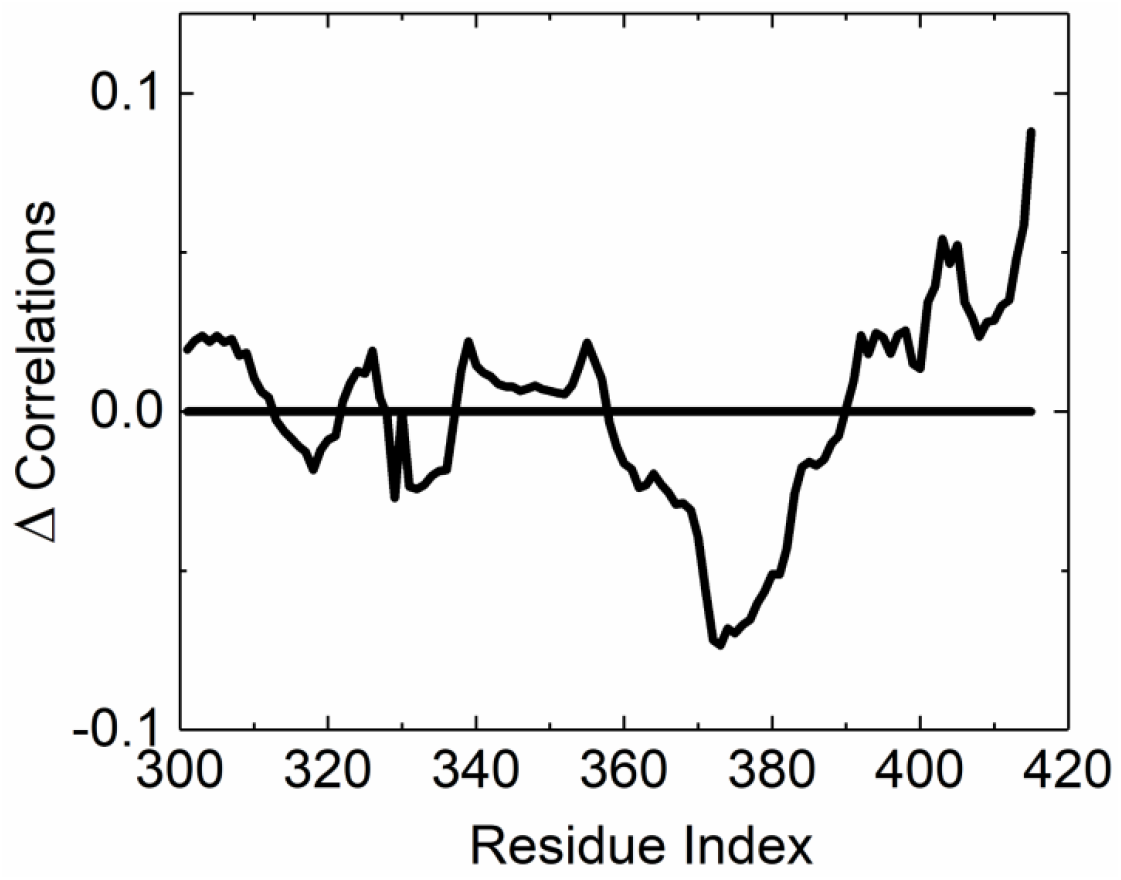
Effect of ligand binding on wild type apo structure. Ordinate values show the differences of correlation of residue 330 with other residues of the protein.

## 4. Conclusion

The all-atom GNM modifies the Kirchoff adjacency matrix of a folded protein by introducing interactions between neighboring heavy atoms. The classical coarse-grained GNM recognized only the alpha carbons and assumed heavy atoms to be phantom-like. The change in the adjacency matrix that comes with the all-atom GNM makes it possible to analyze correlation changes in the system. Our knowledge on the relationship between protein function and correlations of fluctuations has not yet reached an advanced level. A better understanding may be possible by using the all-atom GNM to study the large number of known cases on loss or gain of function upon mutation (or ligand binding) in terms of loss or gain of correlations.

Effect of mutation or ligand binding on distant points in a protein is essentially a problem of allostery. A large volume of research in this field shows that allostery may be divided into two classes, (i) allostery resulting from large conformational changes upon perturbation^24^ and (ii) allostery without conformational change^17^. In case (i), the perturbation changes the adjacency matrix in such a way that new alpha carbon-alpha carbon interactions, i.e., C^α^ -C^α^ terms appear in the Γ matrix. The elements of the new Γ matrix can be obtained only after determining the crystal structure of the perturbed system or after extensive molecular dynamics simulations. The all-atom GNM is most useful for allostery of type (ii) where the atomic structure and conformation of the wild type is sufficient.

Constructing a laboratory mutation and determining the crystal structure of the mutated system appears to be the bottleneck. Therefore, a theoretical model that requires only wild type data is of utmost importance. The approximations involved in the proposed model are not trivial, however. An alternative approach for generating the necessary data and checking the predictions of the all-atom GNM is molecular dynamics simulations^25^.

## Acknowledgment

The author gratefully acknowledges discussions with Professor Turkan Haliloglu.

